# TMvisDB: resource for transmembrane protein annotation and 3D visualization

**DOI:** 10.1101/2022.11.30.518551

**Authors:** Céline Marquet, Anastasia Grekova, Leen Houri, Michael Bernhofer, Luisa F. Jimenez-Soto, Tim Karl, Michael Heinzinger, Christian Dallago, Burkhard Rost

## Abstract

Since the rise of cellular organisms, transmembrane proteins (TMPs) have been crucial to a variety of cellular processes due to their central role as gates and gatekeepers. Despite their importance, experimental high-resolution structures for TMPs remain underrepresented due to technical limitations. With structure prediction methods coming of age, predictions might fill some of the need. However, identifying the membrane regions and topology in three-dimensional structure files requires additional *in silico* prediction. Here, we introduce TMvisDB to sieve through millions of predicted structures for TMPs. This resource enables both, to browse through 46 million predicted TMPs and to visualize those along with their topological annotations. The database was created by joining AlphaFold DB structure predictions and transmembrane topology predictions from the protein language model based method TMbed. We show the utility of TMvisDB for individual proteins through two single use cases, namely the B-lymphocyte antigen CD20 (*Homo sapiens*) and the cellulose synthase (*Novosphingobium sp. P6W*). To demonstrate the value for large scale analyses, we focus on all TMPs predicted for the human proteome. TMvisDB is freely available at tmvis.predictprotein.org.

## Introduction

### Experimental structures for transmembrane proteins remain sparse

Despite many biological processes depending on proteins spanning cell membranes, only 5% of all high-resolution experimental three-dimensional (3D) structures in the Protein Data Bank (PDB) are transmembrane proteins (TMPs) [1]. Although structure-determination methods have advanced significantly, TMPs continue to pose many technical challenges [2]. Therefore, today’s databases often contain only small subsets of membrane proteins, and few focus solely on TMPs or add information on (predicted) membrane embeddings [3–7]. As of November 2022, *mpstruc*, a curated database covers 1,526 unique TMPs with experimental 3D structures [8], which constitutes 1.3% of all PDB structures (114,919 unique sequences redundancy-reduced at 100% identity). Substantially below the estimate of 20-30% of proteins in any proteome being TMPs [9, 10], this imbalance becomes more striking when considering that TMPs make up almost 50% of all drug targets [11].

### Novel resources may push novelty into realm of TMPs

Few computational methods revolutionized a field as quickly and dramatically as AlphaFold2 [12]. Providing reliable 3D predictions, AlphaFold2 and AlphaFold DB (AFDB) are extraordinary catalysts for research in artificial intelligence (AI) and the life sciences [13]. Similarly to structure predictors advancing rapidly [14–17], methods to quickly and reliably search and cluster the exploding sequence space (>2.4 billion in metagenomic databases) are continuously improving [18–20]. An orthogonal approach to leverage the wealth of sequence data was introduced by protein language models (pLMs) [21, 22]. Such pLMs learn from protein sequences without experimental annotations (dubbed ‘unlabeled’ in AI jargon) to extract meaningful numeric representations (embeddings) encoding sequences as vectors. The knowledge acquired during pre-training on raw sequences can readily be transferred to downstream predictions of per-residue [10, 23–26], and per-protein phenotypes [27–30]. Similar concepts successfully generate new sequences in the context of protein design [31–33]. The unparalleled number of available sequences, pLM embeddings, and experimental and predicted structures, may help to overcome limitations for studying TMPs *in vitro* and *in vivo*. Combining the aforementioned *in silico* methods and resources enables discovering previously unknown TMPs and exploring both known and unknown TMPs in exceptional detail. An example for the usefulness of TMP predictions is the *TmAlphaFold database*, taking AlphaFold2 structures as input to provide membrane localization for almost 216 thousand predicted alpha-helical TMPs [34].

Here, we introduce TMvisDB, a collection of over 46 million (46M) predicted alpha-helical and beta-barrel TMPs with their AlphaFold2-predicted 3D structure as well as membrane topology and annotation predicted by the pLM-based method TMbed [10]. TMvisDB was created by applying TMbed for all proteins in AFDB (>200M, 07/22). TMbed is either *on par* with or outperforms all other TMP prediction methods in classifying residues as transmembrane alpha-helix (TMH) or transmembrane beta-strand (TMB). TMbed also predicts the orientation of TM-segments with respect to the membrane (dubbed *topology*), and signal peptides (SP). The latter reaches the binary classification performance of SignalP6, without distinguishing different SP types [10, 35]. TMvisDB is the first resource to gather such a large number of per-residue transmembrane topology annotations and, compared to existing TMP databases [8, 34], expanding the exploration space of easily browsable TMPs by two to four orders of magnitude. Providing interactive visualizations of TMP annotations with their respective AlphaFold2 structures, the intuitive interface of tmvis.predictprotein.org enables novices and experts in fields such as microbiology, drug development and bioinformatics to investigate small- and large-scale hypotheses on TMPs.

## Results

### Over 46M predicted transmembrane proteins

TMvisDB is a non-relational database, with 46,048,450 proteins (12/22, Table 1), comprising 23% of AlphaFold DB (AFDB, >200M, 07/22) [13]. Transmembrane proteins (TMPs) and their topology were predicted for proteins in AFDB using the protein language model based, state-of-the-art TMP prediction method TMbed [10]. About 95.3% of the proteins in TMvisDB have predicted transmembrane alpha-helices (TMHs), 4.6% contain predicted transmembrane beta-strands (TMBs), and 0.1% contain both, likely providing a lower estimate for the error of our approach while no convincing current evidence supports or rules out the existence of such proteins. Signal peptides (SP) were predicted by TMbed for 13.6% of all TMPs (archaea: 13.83%, bacteria: 14.5%, eukaryotes: 15.3%), compared to 7.1% of all proteins in UniProt with AlphaFold2 structures and transmembrane annotation.

**Table 1:**
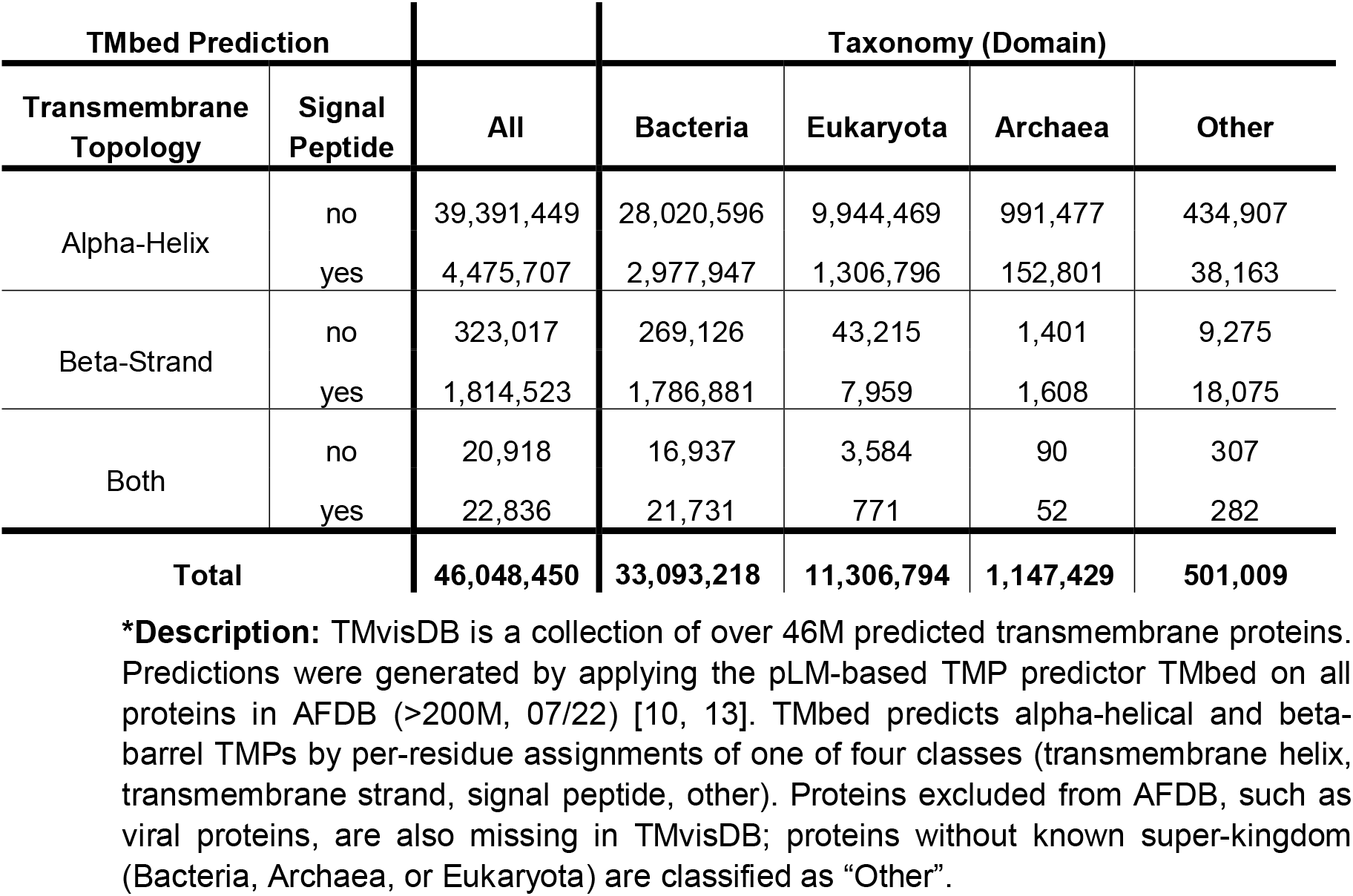
Summary of proteins in TMvisDB*.

### Intuitive browsing and visualization

The website tmvis.predictprotein.org enables interactive access to browse and visualize all TMPs in TMvisDB. For large-scale analyses, the database can be filtered by taxonomy (UniProt organism identifier, domain, kingdom), predicted topology (occurrence of TMH/TMB/both), protein length, and presence/absence of predicted SP (Fig. 1 left). Each database record contains the UniProt accession number (UA, unique identifier), taxonomy information parsed from UniProt, protein length, protein sequence (one-letter amino acid code), and per-residue TMbed predictions (SP: signal peptide, TMH: transmembrane alpha-helix, TMB: transmembrane beta-strand, other; Fig. 1 right), and results are downloadable in csv-format (comma-separated columns in ASCII). Individual TMPs are selectable by their UA to visualize the corresponding AlphaFold2 3D structure, where each residue can be colored by (1) one of seven topology states (inside-to-outside/outside-to-inside TMH/TMB, intra-/extracellular residues, SP), or (2) the AlphaFold2 per-residue confidence metric (the so called predicted local distance difference test - pLDDT; Fig. 2, Supporting Online Material (SOM) Fig.S1-S3). Protein sequence annotations are shown below the 3D visualization along with links to UniProt [36], LambdaPP [37], a webserver providing further protein-specific phenotype predictions, and Foldseek [38] for fast and highly sensitive 3D structure search and alignment.

**Fig. 1:**
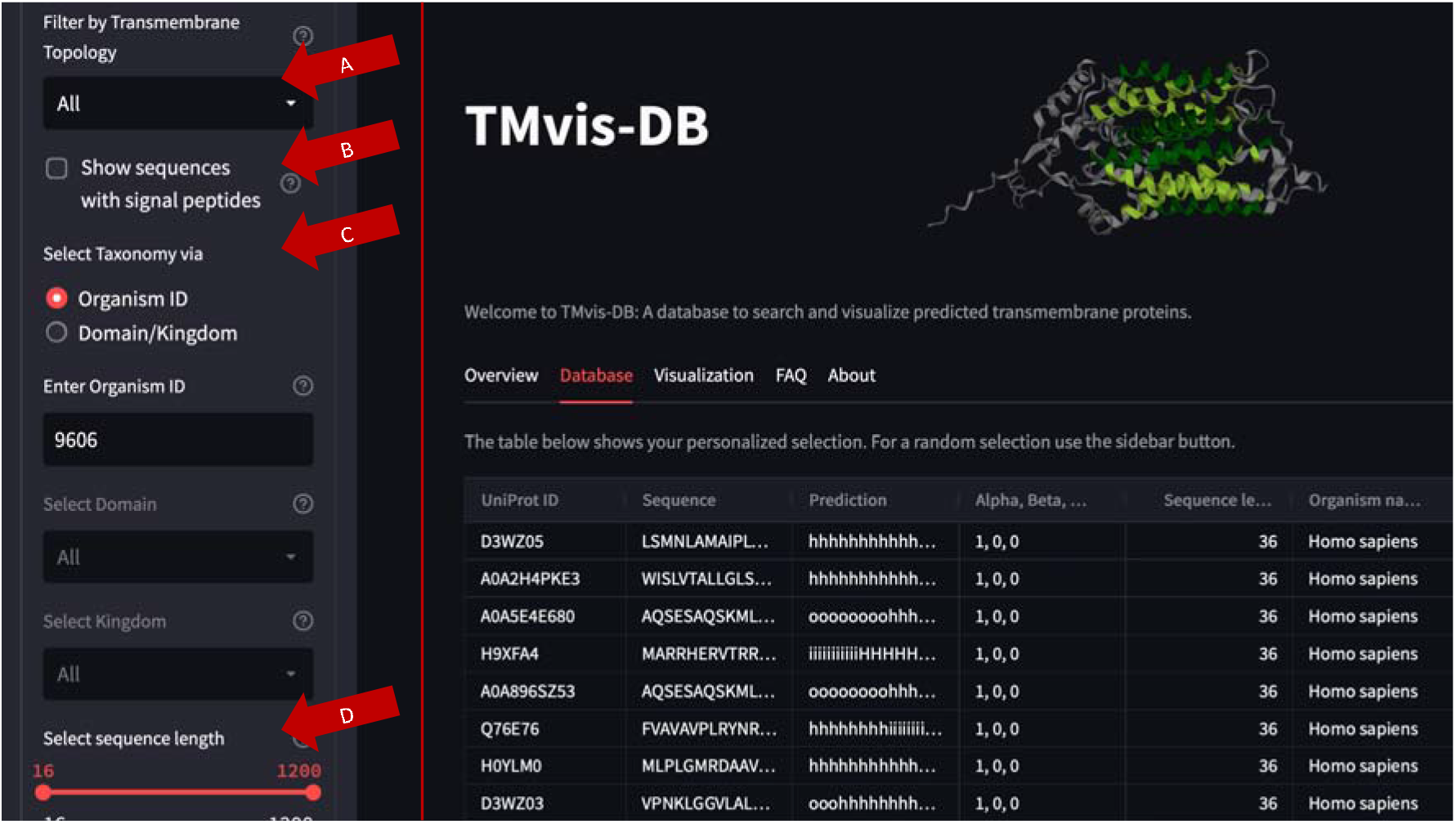
TMvisDB interface. **Left** (light gray): The side bar shows available filters for browsing TMvisDB. Currently, users can filter by (A) Transmembrane topology (alpha-helix, beta-strand), (B) Signal Peptides (SP) (include/exclude sequences with predicted SP), (C) Taxonomy (UniProt Organism Identifier, Domain, Kingdom), and (D) Sequence length. **Right** (dark gray): Partial output of *homo sapiens* TMvisDB selection (UniProt organism identifier 9606) with record features: UniProt accession, sequence, per-residue TMbed predictions [10], sequence classification (alpha-helix, beta-strand, SP), sequence length, organism name, UniProt organism identifier, and lineage.

**Fig. 2:**
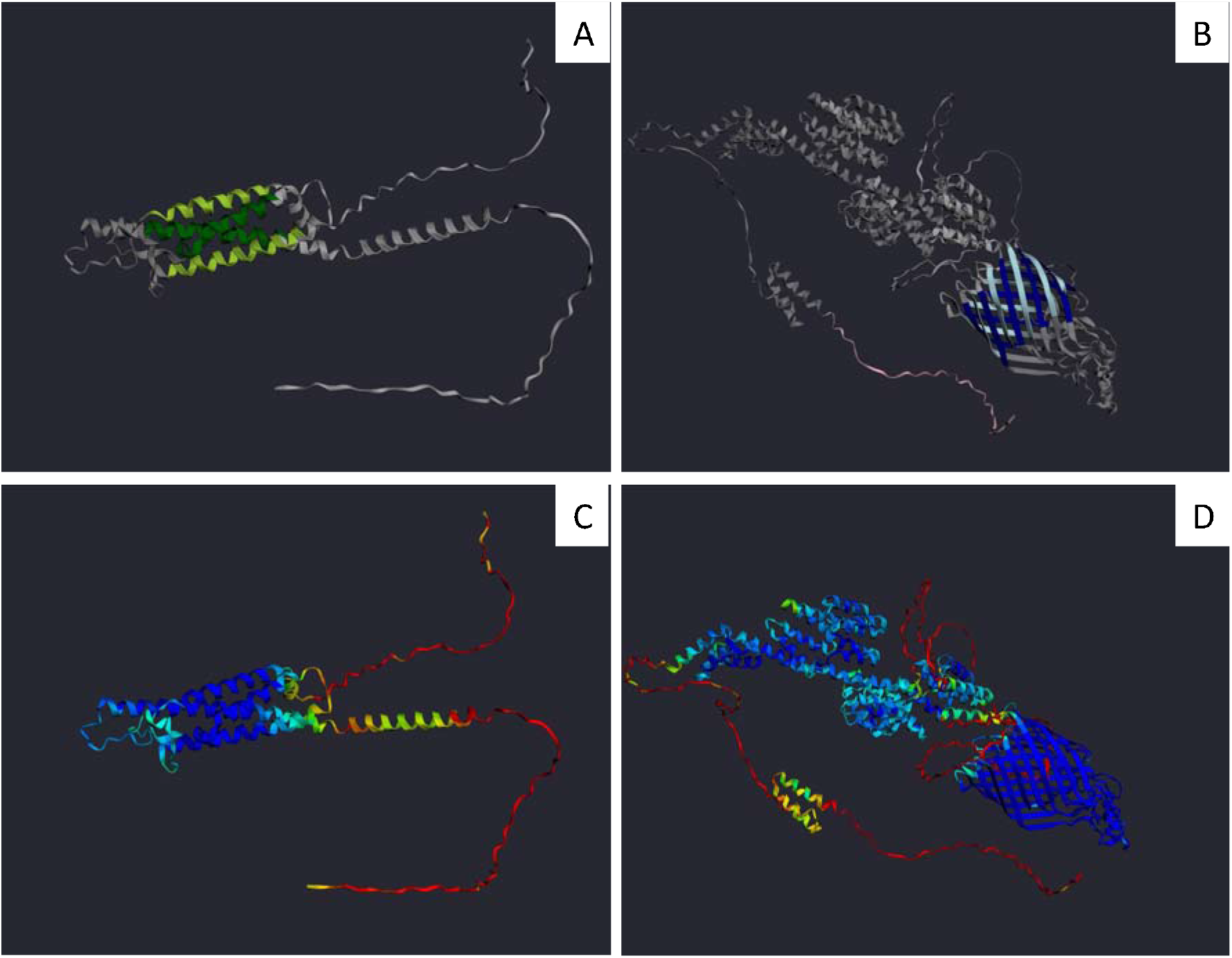
TMvisDB single protein use cases. Two proteins, B-lymphocyte antigen CD20 (P11836, *Homo sapiens*) and cellulose synthase protein (A0A2Z5E6S1, *Novosphingobium sp. P6W*) were chosen for detailed analysis. AlphaFold2 structure predictions for CD20 are shown in the left two panels **(A&C)** and predictions for the cellulose synthase protein are shown in the two right panels **(B&D)** [12]. The upper row **(A&B)** colors per-residue transmembrane topology predicted by TMbed according to the following color-scheme: inside-to-outside alpha-helix (light green), outside-to-inside alpha-helix (dark green), inside-to-outside beta-strand (light blue), outside-to-inside beta-strand (dark blue), signal peptide (pink), other (grey). The structures in the panels of the lower row **(C&D)** are colored by the AlphaFold2 per-residue confidence metric, predicted local distance difference test (pLDDT), according to the following color-scheme: very low pLDDT ≤ 50 (red), low 50 < pLDDT ≤ 70 (yellow), confident 70 < pLDDT ≤ 90 (green), very confident pLDDT > 90 (blue). For both proteins, the predicted transmembrane topology aligns with the predicted structure, and with regions of high pLDDT in a sense that transmembrane residues are predicted with high confidence while loopy non-membrane residues have a higher uncertainty according to pLDDT.

### Use case 1: B-lymphocyte antigen CD20 - expanding annotations

To demonstrate the TMvisDB web interface for a single protein, we selected the *B-lymphocyte antigen CD20* (UA: P11836, Fig. 2A,C) following the suggestion of a biochemist and because, as in many cases, reliable predictions might help answering some of the open questions about its molecular and biological function. Although a well-studied TMP and widespread therapeutical target, no experimental high-resolution structure is available for CD20 in isolation. Recently, it has been demonstrated that the loss of CD20 disrupts membrane organization, and that it acts as gatekeeper of a membrane nanodomain and the resting state of naive B-cells [39]. CD20 has been suggested to form a homo-oligomer, and so far, its experimental structure has only been determined in complexes [40, 41]. The TMbed prediction suggests four TMHs (length 18-24 residues): (1) inside-to-outside residues (residues 51-72), followed by eight outside residues, (2) outside-to-inside residues (residues 81-99), followed by fifteen inside residues, (3) inside-to-outside (residues 115-139), followed by 48 outside residues, and (4) outside-to-inside (residues 188-211). Neither TMbed, nor SignalP6.0 predict a SP. The TMHs predicted by TMbed fit to the structure predicted by AlphaFold2 with high confidence (Fig. 2C). While TMvisDB annotations deviate fundamentally from the transmembrane regions in UniProt [42], they mostly agree with and confirm the TMHs and orientation seen in the experimental structure [40]. Furthermore, the TMvisDB transmembrane topology and structure predictions support studies suggesting CD20 to be a member of the MS4A family of TMPs, forming a non-glycosylated tetraspanin with one shorter and one longer extracellular loop, and both C- and N-termini inside the cytosol (Fig. 2A) [43, 44].

### Use case 2: Cellulose synthase - exploring novel TMP annotation

TMvisDB facilitates the analysis of previously unstudied TMPs, such as the cellulose synthase protein (UA: A0A2Z5E6S1) of *Novosphingobium sp. P6W*, a gram-negative bacterium (Fig. 2 B,D). According to automatic annotation for unreviewed records by UniProt, it lies in the prokaryotic outer membrane, and is associated with glycan metabolism and bacterial cellulose biosynthesis [45]. While automatic UniProt annotation shows 29 residues as SP, TMbed predicted 40 residues. This longer prediction is in accordance with a strong prediction by the expert method SignalP6.0 (cleavage site at 40, probability 0.97) [35]. Beginning at residue 820, TMbed predicted twelve TMB, which align with the beta-barrel predicted by AlphaFold2 (Fig. 2B). The high pLDDT of the transmembrane residues added evidence confirming the prediction of an unknown TMB (Fig. 2D). Minor deviations remain as some predicted TMBs were slightly shorter than the AlphaFold2 barrel structure (Fig. 2B, lower right). We could find no conclusive evidence whether the prediction of a shorter TMB was due to the natural embedding in the membrane, or to a few residues being predicted by TMbed as false negatives, or to protein dynamics triggered upon binding that might lead AlphaFold2 to predict a slightly altered state of the protein.

### Use case 3: *Homo sapiens* - combining large- and small-scale analyses

In addition to studies of individual proteins, TMvisDB enables large-scale analyses. TMvisDB contains 20.7% of the human proteins in AFDB (07/22), accessible via the human UniProt organism identifier (no. 9606). Out of this set, we investigated the only proteins for which TMbed predicted both TMH and TMB, seemingly being outliers (Deoxyribonuclease I fragment (UA: A0A192ZHB2; Fig. S1), two sequence-similar perforins (UA: B3KUT2, Q2M385; Fig. S2), three sequence-similar DnaJ homologues (UA: B4DGD5, B4DPK2, Q9NVH1; Fig. S3)).

For the Deoxyribonuclease I fragment, TMbed predicted one inside-to-outside TMH and one outside-to-inside TMB; AlphaFold2 predicted an alpha-helix and beta-strand in the same region with only the beta-strand predicted confidently and without any loop/bend between the two predicted secondary structure segments (Fig. S1). For this fragmented protein, these inconsistencies suggested that either one or both predictions were at best partially correct.

Whether or not the topology prediction for the perforins was correct also remained unclear. The TMbed-predicted TMH aligned with a helix predicted by AlphaFold2 (Fig. S2), the transmembrane region automatically annotated by UniProt [46], and research suggesting that perforins are able to insert in target cellular membranes [47]. While for both the short predicted TMB (5-6 residues) neither aligns with AlphaFold2 nor with UniProt, the Macrophage-expressed gene 1 protein (UA: Q2M385) is thought to homo-oligomerize into a pore-forming ring comprising a large beta-barrel [48]. To investigate this further, we analyzed other perforin-like proteins targeting similar membranes (*Toxoplasma gondii* (G3G7T0, intracellular parasite in white blood cells), *Plasmodium falciparum* (Q9U0J9, intracellular parasite in red blood cells)), none of which showed similar predictions. Based on this and on the fact that predictions from AlphaFold2 and UniProt provided no support for the TMB-hypothesis, we took this to be an incorrect prediction.

For the final case of TMbed predicting TMH and TMB, namely the DnaJ homologues, the AlphaFold2 structure prediction appeared to confirm TMbed; the length of the TMH and the TMB agreed with the TMbed-predicted membrane boundaries (Fig. S3). Experimental research describes DnaJ as part of the mitochondrial intermembrane space bridging (MIB) complex spanning both mitochondrial membranes, and as associated with the mitochondrial contact site and cristae junction organizing system (MICOS) [49, 50]. Although there is not yet experimental evidence of the two transmembrane domains from DnaJ playing a role in its function, the experimental existing data associating DnaJ proteins with the MICOS assembly in mitochondria and its possible role in cristae organization could explain the use of two different transmembrane domains. Thus, this might be a new discovery or might await experimental falsification.

## Discussion

### Organizing the growing number of available structures

The rapid growth available protein sequences and 3D structures (experimental and computational) increases the need to group specific proteins into easily accessible collections. Protein language models (pLMs) allow scanning large sequence databases for certain functionalities at unprecedented speed, regardless of sequence alignment availability [37, 51] and even without considering sequence similarity [38]. This development allowed to apply the pLM-based method TMbed on most proteins in UniProt and extract a subset containing only predicted transmembrane proteins (TMPs). The database and web application TMvisDB offered a straightforward search functionality and visualization interface. While several accurate structure prediction methods have been made available over the last year [14–17], we chose to enhance TMvisDB sequence annotations with AlphaFold2 [12] predictions that have been shown to perform well in structural analysis of transmembrane proteins (TMPs) [52], and have, therefore, been successfully applied as input by resources such as the *TmAlphaFold database* that collects alpha-helical TMPs [34].

### Benefits and limitations often hand in hand

To complement current resources and help facilitate a better understanding of the molecular and biological role of TMPs, TMvisDB combines predicted per-residue transmembrane (TM) and signal peptide annotations with 3D protein structure. The resources employed by TMvisDB rely on different input, i.e., the pLM-based method TMbed takes single sequences as input whereas AlphaFold2 requires multiple sequence alignments. Hence, the one-dimensional topology prediction (TMbed) and the three-dimensional structure prediction (AlphaFold2) complement each other and provide independent evidence. This leads to findings such as the cellulose synthase protein, where predictions align but it remains unclear to which extent the beta-barrel is embedded in the membrane. The intuitive color-coding for TMbed transmembrane topology or AlphaFold2 confidence enhances critical assessment. As shown by the *B-lymphocyte antigen CD20*, TMvisDB can support *in silico* confirmation of TM-regions and their orientations, which in turn allows a higher rate of success for the design of experimental molecular studies. Yet, the monomeric structure of CD20, while displayed with seemingly correct TM-topology, falls short to allow detailed analysis of a protein likely forming homo-oligomeric complexes. Until tools such as AlphaFold-Multimer become available on a larger scale [53], users have to investigate single-use cases.

Despite the outstandingly low false positive rate of TMbed (less than 1% for both types of TMPs), at least some of the proteins predicted with both TM alpha-helix (TMH) and TM beta-strand (TMB) must fall into this category as no such mixed protein has been observed before. TMvisDB enables the evaluation of such cases by providing access to quick and easy comparative studies as shown for the perforins from *homo sapiens*. The comparison with perforin-like proteins in other organisms allows identifying false positives. Nevertheless, this use case also revealed some challenges. Over 38 thousand (38K) proteins in TMvisDB are human TMPs, which seems somewhat nonsensical, given there are likely no more than 20K human proteins [54]. This surprising number originates from the *“logic”* of our pipeline: TMvisDB is based on AlphaFold DB (AFDB), which currently holds records for 186k human proteins, i.e., almost an order of magnitude more than exist. AFDB includes unreviewed proteins, fragments such as the Deoxyribonuclease I fragment, and it does not account for redundancy. While having structures even for redundant sequences is mostly beneficial, the scale of TMvisDB makes searchability and visualization challenging. For more efficient filtering, we will add options to filter features such as Gene Ontology (GO) annotations and binding [25, 30]. We also intend to speed-up TMvisDB and to update it regularly with new AFDB releases. With the help of user feedback, we hope to provide a highly valuable tool for TMP research.

## Materials and Methods

### TMbed transmembrane topology prediction

TMbed predicts alpha-helical and beta-barrel transmembrane proteins (TMPs) by assigning one of four classes (transmembrane alpha-helix (TMH), transmembrane beta-strand (TMB), signal peptide (SP), or other) to each residue of a protein sequence [10]. For TMH and TMB segments it also predicts their orientation within the membrane, i.e., whether the segment is from inside-to-outside or outside-to-inside. However, focusing on transmembrane segments, it does not predict re-entrant regions. Performing on par or better than state-of-the-art methods, TMbed correctly identifies 98±1% of alpha-helical and 94±8% of beta-barrel TMPs, while maintaining a false positive rate (i.e., predicting non-TMPs as TMPs) of less than 1% for both types of TMPs. On average, it places about 9 of 10 predicted TMHs and TMBs within five residues of their correct location in the protein sequence. Requiring only protein language model representations (embeddings) from single amino acid sequences as input, TMbed can predict UniProtKB/Swiss-Prot (566,976 sequences) in less than 9 hours, making it well suited for high-throughput applications such as TMvisDB.

### TMvis database

The TMvisDB is a NO-SQL JSON-based database implemented with MongoDB. Each database entry is identifiable via its unique UniProt accession (UA), and contains fields describing protein and sequence, prediction and annotation, and taxonomic information. Further, if available, it contains annotations of experimentally derived topology information of TOPDB [55], and single alpha-helix transmembrane proteins of Membranome [3]. TMvisDB contains all proteins of AlphaFold DB (AFDB) [13], which have transmembrane components predicted by the method TMbed. As of January ‘23, AFDB has the following sequence coverage limitations (which in turn also apply to TMvisDB): sequences are (1) inside length range of minimum 16 amino acids, maximum of 2,700 for Swiss-Prot, and 1,280 for all other; (2) not containing non-standard amino acids; (3) in the UniProt reference proteome “one sequence per gene” FASTA file; (4) not added or modified by UniProt in more recent release; (5) not a viral protein. TMvisDB is hosted at Rostlab (https://rostlab.org).

### TMvis visualization

TMvis combines AlphaFold2 structures from AFDB with predicted TMP annotations into interactive 3D visualizations of protein structures embedded into membranes using py3Dmol [56]. To visualize proteins with TMvis locally, the user must download the respective proteins from AFDB and run TMbed predictions, either via its repository (https://github.com/Rostlab/tmbed), bioembeddings [51], or LambdaPP [37]. The predictions can then be visualized with TMvis (https://github.com/Rostlab/TMvis).

### TMvis web application

The TMvisDB web app is built with the python-based open-source library Streamlit (https://streamlit.io/) that enables customized web apps for machine learning and data science. The TMvisDB web app is hosted on the Streamlit cloud, and accessible via tmvis.predictprotein.org. It connects to TMvisDB at Rostlab via PyMongo to enable browsing the 46 million predicted transmembrane proteins. The user can make selections with the following filters: transmembrane topology (TMH, TMB), include/exclude sequences with predicted SP, taxonomy (UniProt organism identifier, domain, kingdom), and protein length (Fig. 1). All results are downloadable via the web app. Further, single proteins of TMvisDB can be selected for 3D visualization of per-residue transmembrane topology annotation (Fig. 2). A table below the 3D visualization allows to compare TMbed predictions [10] to transmembrane topology information (if available) of TOPDB [55], Membranome [3], UniProt [36], and the TmAlphaFold database [34]. Proteins can either be selected from the table generated while browsing TMvisDB, or via the UniProt accession number. The AlphaFold2 structures of a protein is fetched from AFDB, and then shown with the corresponding color code of the predicted topology or the AlphaFold2 pLDDT score. The visualization is based on TMvis (https://github.com/Rostlab/TMvis) and implemented using Stmol [57]. The web app source code is accessible via github.com/Rostlab/TMvisDB.

## Abbreviations

AFDB: AlphaFold DB
AI: artificial intelligence
ASCII: American Standard Code for Information Interchange
GO: Gene Ontology
MIB: mitochondrial intermembrane space bridging complex
MICOS: mitochondrial contact site and cristae junction organizing system
PDB: Protein Data Bank
pLDDT: predicted local distance difference test
pLM: protein language model
SAM: mitochondrial outer membrane sorting assembly machinery
SOM: supporting online material
SP: signal peptide
TM: transmembrane
TMB: transmembrane beta strand
TMH: transmembrane alpha helix
TMP: transmembrane protein
TOPDB: Topology Database of Transmembrane Proteins
UA: UniProt accession number
3D: three dimensional

## Accession Numbers

UniProt accessions: A0A2Z5E6S1, P11836, A0A192ZHB2, B3KUT2, Q2M385, B4DGD5, B4DPK2, Q9NVH1, G3G7T0, Q9U0J9

UniProt organism identifier: 9606

## Acknowledgements

Thanks to Inga Weise for support with many aspects of this work, and to Arne Skerra for suggesting the CD20 use case. Thanks to John Jumper and his team for the breakthrough development of AlphaFold2 and making code and predictions freely available. Further, thanks to all contributing to open-source programming libraries. Last, not least, thanks to all who deposit experimental data in public databases, to those who maintain these databases, and those who make methods available enriching experimental data.

## Author contributions

C.M. co-conceived project, advised development of visualization pipeline TMvis, generated and evaluated TMvisDB, implemented website, wrote manuscript. A.G. developed TMvis, contributed to early case studies. L.H. contributed to case studies. M.B. developed TMbed, advised project and manuscript. L.J.-S. advised and contributed to writing case studies. T.K. built and managed deployment of publicly available database server hosting TMvisDB, provided beneficial feedback. M.H. co-conceived project, implemented pipeline to generate TMbed predictions for AlphaFold DB. C.D. contributed advice on project conception and implementation. B.R. supervised and guided the work.

## References

[1] Burley SK, Bhikadiya C, Bi C, Bittrich S, Chen L, Crichlow GV, et al. RCSB Protein Data Bank: powerful new tools for exploring 3D structures of biological macromolecules for basic and applied research and education in fundamental biology, biomedicine, biotechnology, bioengineering and energy sciences. Nucleic Acids Research. 2021;49:D437–D51.

[2] Li F, Egea PF, Vecchio AJ, Asial I, Gupta M, Paulino J, et al. Highlighting membrane protein structure and function: A celebration of the Protein Data Bank. Journal of Biological Chemistry. 2021;296.

[3] Lomize AL, Schnitzer KA, Todd SC, Cherepanov S, Outeiral C, Deane CM, et al. Membranome 3.0: Database of single-pass membrane proteins with AlphaFold models. Protein Science. 2022;31:e4318.

[4] Lomize MA, Pogozheva ID, Joo H, Mosberg HI, Lomize AL. OPM database and PPM web server: resources for positioning of proteins in membranes. Nucleic Acids Research. 2012;40:D370–6.

[5] Postic G, Ghouzam Y, Etchebest C, Gelly J-C. TMPL: a database of experimental and theoretical transmembrane protein models positioned in the lipid bilayer. Database. 2017;2017:bax022.

[6] Newport TD, Sansom MS P, Stansfeld PJ. The MemProtMD database: a resource for membrane-embedded protein structures and their lipid interactions. Nucleic Acids Research. 2019;47:D390–D7.

[7] Tordai H, Suhajda E, Sillitoe I, Nair S, Varadi M, Hegedus T. Comprehensive Collection and Prediction of ABC Transmembrane Protein Structures in the AI Era of Structural Biology. International Journal of Molecular Sciences. 2022;23:8877.

[8] White SH. mpstruc: Membrane Proteins of Known Structure. 1998.

[9] Fagerberg L, Jonasson K, von Heijne G, Uhlén M, Berglund L. Prediction of the human membrane proteome. Proteomics. 2010;10:1141–9.

[10] Bernhofer M, Rost B. TMbed: transmembrane proteins predicted through language model embeddings. BMC Bioinformatics. 2022;23:326.

[11] Santos R, Ursu O, Gaulton A, Bento AP, Donadi RS, Bologa CG, et al. A comprehensive map of molecular drug targets. Nat Rev Drug Discov. 2017;16:19–34.

[12] Jumper J, Evans R, Pritzel A, Green T, Figurnov M, Ronneberger O, et al. Highly accurate protein structure prediction with AlphaFold. Nature. 2021;596:583–9.

[13] Varadi M, Anyango S, Deshpande M, Nair S, Natassia C, Yordanova G, et al. AlphaFold Protein Structure Database: massively expanding the structural coverage of protein-sequence space with high-accuracy models. Nucleic Acids Research. 2022;50:D439–D44.

[14] Ahdritz G, Bouatta N, Kadyan S, Xia Q, Gerecke W, AlQuraishi M. OpenFold. 2021.

[15] Baek M, Baker D. Deep learning and protein structure modeling. Nat Methods. 2022;19:13–4.

[16] Lee JH, Yadollahpour P, Watkins A, Frey NC, Leaver-Fay A, Ra S, et al. EquiFold: Protein Structure Prediction with a Novel Coarse-Grained Structure Representation. bioRxiv; 2022.

[17] Mirdita M, Schütze K, Moriwaki Y, Heo L, Ovchinnikov S, Steinegger M. ColabFold: making protein folding accessible to all. Nat Methods. 2022;19:679–82.

[18] Mirdita M, Steinegger M, Söding J. MMseqs2 desktop and local web server app for fast, interactive sequence searches. Bioinformatics. 2019;35:2856–8.

[19] Buchfink B, Reuter K, Drost H-G. Sensitive protein alignments at tree-of-life scale using DIAMOND. Nat Methods. 2021;18:366–8.

[20] Steinegger M, Mirdita M, Soding J. Protein-level assembly increases protein sequence recovery from metagenomic samples manyfold. Nat Methods. 2019;16:603–6.

[21] Rives A, Meier J, Sercu T, Goyal S, Lin Z, Liu J, et al. Biological structure and function emerge from scaling unsupervised learning to 250 million protein sequences. Proceedings of the National Academy of Sciences. 2021;118:e2016239118.

[22] Elnaggar A, Heinzinger M, Dallago C, Rehawi G, Wang Y, Jones L, et al. ProtTrans: Toward Understanding the Language of Life Through Self-Supervised Learning. IEEE Transactions on Pattern Analysis and Machine Intelligence. 2022;44:7112–27.

[23] Marquet C, Heinzinger M, Olenyi T, Dallago C, Erckert K, Bernhofer M, et al. Embeddings from protein language models predict conservation and variant effects. Hum Genet. 2022;141:1629–47.

[24] Ilzhoefer D, Heinzinger M, Rost B. SETH predicts nuances of residue disorder from protein embeddings. Front Bioinform. 2022.

[25] Littmann M, Heinzinger M, Dallago C, Weissenow K, Rost B. Protein embeddings and deep learning predict binding residues for various ligand types. Sci Rep. 2021;11:23916.

[26] Meier J, Rao R, Verkuil R, Liu J, Sercu T, Rives A. Language models enable zero-shot prediction of the effects of mutations on protein function. bioRxiv. 2021:2021.07.09.450648.

[27] Bileschi ML, Belanger D, Bryant DH, Sanderson T, Carter B, Sculley D, et al. Using deep learning to annotate the protein universe. Nat Biotechnol. 2022;40:932–7.

[28] Hamid M-N, Friedberg I. Identifying antimicrobial peptides using word embedding with deep recurrent neural networks. Bioinformatics. 2019;35:2009–16.

[29] Stärk H, Dallago C, Heinzinger M, Rost B. Light attention predicts protein location from the language of life. Bioinformatics Advances. 2021;1:vbab035.

[30] Littmann M, Heinzinger M, Dallago C, Olenyi T, Rost B. Embeddings from deep learning transfer GO annotations beyond homology. Sci Rep. 2021;11:1160.

[31] Ferruz N, Schmidt S, Höcker B. ProtGPT2 is a deep unsupervised language model for protein design. Nat Commun. 2022;13:4348.

[32] Nijkamp E, Ruffolo J, Weinstein EN, Naik N, Madani A. ProGen2: Exploring the Boundaries of Protein Language Models. arXiv; 2022.

[33] Moffat L, Kandathil SM, Jones DT. Design in the DARK: Learning Deep Generative Models for De Novo Protein Design. bioRxiv; 2022.

[34] Dobson L, Szekeres LI, Gerdán C, Langó T, Zeke A, Tusnády GE. TmAlphaFold database: membrane localization and evaluation of AlphaFold2 predicted alpha-helical transmembrane protein structures. Nucleic Acids Research. 2022:gkac928.

[35] Teufel F, Almagro Armenteros JJ, Johansen AR, Gíslason MH, Pihl SI, Tsirigos KD, et al. SignalP 6.0 predicts all five types of signal peptides using protein language models. Nat Biotechnol. 2022;40:1023–5.

[36] The UniProt C. UniProt: the universal protein knowledgebase in 2021. Nucleic Acids Research. 2021;49:D480–D9.

[37] Olenyi T, Marquet C, Heinzinger M, Kröger B, Nikolova T, Bernhofer M, et al. LambdaPP: Fast and accessible protein-specific phenotype predictions. bioRxiv; 2022.

[38] Kempen Mv, Kim SS, Tumescheit C, Mirdita M, Gilchrist CLM, Söding J, et al. Foldseek: fast and accurate protein structure search. bioRxiv; 2022.

[39] Kläsener K, Jellusova J, Andrieux G, Salzer U, Böhler C, Steiner SN, et al. CD20 as a gatekeeper of the resting state of human B cells. Proceedings of the National Academy of Sciences. 2021;118:e2021342118.

[40] Rougé L, Chiang N, Steffek M, Kugel C, Croll TI, Tam C, et al. Structure of CD20 in complex with the therapeutic monoclonal antibody rituximab. Science. 2020;367:1224–30.

[41] Kumar A, Planchais C, Fronzes R, Mouquet H, Reyes N. Binding mechanisms of therapeutic antibodies to human CD20. Science. 2020;369:793–9.

[42] UniProtKB. B-lymphocyte antigen CD20 - Homo sapiens | UniProtKB | UniProt. 2022.

[43] Oettgen HC, Bayard PJ, Van Ewijk W, Nadler LM, Terhorst CP. Further Biochemical Studies of the Human B-Cell Differentiation Antigens B1 and B2. Hybridoma. 1983;2:17–28.

[44] Bubien JK, Zhou LJ, Bell PD, Frizzell RA, Tedder TF. Transfection of the CD20 cell surface molecule into ectopic cell types generates a Ca2+ conductance found constitutively in B lymphocytes. Journal of Cell Biology. 1993;121:1121–32.

[45] UniProtKB. Cellulose synthase - Novosphingobium sp. P6W | UniProtKB | UniProt. 2022.

[46] UniProtKB. MPEG1 - Macrophage-expressed gene 1 protein - Homo sapiens (Human) | UniProtKB | UniProt. 2022.

[47] Osińska I, Popko K, Demkow U. Perforin: an important player in immune response. Cent Eur J Immunol. 2014;39:109–15.

[48] Pang SS, Bayly-Jones C, Radjainia M, Spicer BA, Law RHP, Hodel AW, et al. The cryoEM structure of the acid activatable pore-forming immune effector Macrophage-expressed gene 1. Nat Commun. 2019;10:4288.

[49] Xie J, Marusich MF, Souda P, Whitelegge J, Capaldi RA. The mitochondrial inner membrane protein mitofilin exists as a complex with SAM50, metaxins 1 and 2, coiled-coil-helix coiled-coil-helix domain-containing protein 3 and 6 and DnaJC11. FEBS Lett. 2007;581:3545–9.

[50] Huynen MA, Mühlmeister M, Gotthardt K, Guerrero-Castillo S, Brandt U. Evolution and structural organization of the mitochondrial contact site (MICOS) complex and the mitochondrial intermembrane space bridging (MIB) complex. Biochimica et Biophysica Acta (BBA) - Molecular Cell Research. 2016;1863:91–101.

[51] Dallago C, Schütze K, Heinzinger M, Olenyi T, Littmann M, Lu AX, et al. Learned Embeddings from Deep Learning to Visualize and Predict Protein Sets. Current Protocols. 2021;1:e113.

[52] Hegedüs T, Geisler M, Lukács GL, Farkas B. Ins and outs of AlphaFold2 transmembrane protein structure predictions. Cell Mol Life Sci. 2022;79:73.

[53] Evans R, O’Neill M, Pritzel A, Antropova N, Senior A, Green T, et al. Protein complex prediction with AlphaFold-Multimer. bioRxiv; 2022.

[54] Wilhelm M, Schlegl J, Hahne H, Gholami AM, Lieberenz M, Savitski MM, et al. Mass-spectrometry-based draft of the human proteome. Nature. 2014;509:582–7.

[55] Dobson L, Lango T, Remenyi I, Tusnady GE. Expediting topology data gathering for the TOPDB database. Nucleic Acids Res. 2015;43:D283–9.

[56] Rego N, Koes D. 3Dmol.js: molecular visualization with WebGL. Bioinformatics (Oxford, England). 2015;31:1322–4.

[57] Nápoles-Duarte JM, Biswas A, Parker MI, Palomares-Baez JP, Chávez-Rojo MA, Rodríguez-Valdez LM. Stmol: A component for building interactive molecular visualizations within streamlit web-applications. Front Mol Biosci. 2022;9:990846.

