## Supporting Online Material for "TMvisDB: resource for transmembrane protein annotation and 3D visualization"

Supporting Online Material (SOM) for:
TMvisDB: resource for transmembrane protein annotation and 3D visualization

Céline Marquet*^1,2^, Anastasia Grekova^1^, Leen Houri^3,4^, Michael Bernhofer^1,2,5^, Luisa F. Jimenez-Soto^6^, Tim Karl^1^, Michael Heinzinger^1,2^, Christian Dallago^1,7,8^, Burkhard Rost^1,3,9^

1	TUM (Technical University of Munich) Department of Informatics, Bioinformatics & Computational Biology - i12, Boltzmannstr. 3, 85748 Garching/Munich, Germany

2	TUM Graduate School, Center of Doctoral Studies in Informatics and its Applications (CeDoSIA), Boltzmannstr. 11, 85748 Garching, Germany

3	TUM School of Life Sciences Weihenstephan (WZW), Alte Akademie 8, Freising, Germany

4	DTU, Technical University of Denmark, Department of Biotechnology and Biomedicine, Søltofts Plads, Building 221, DK-2800 Kgs. Lyngby, Denmark

5	BASF SE, Carl-Bosch-Str. 38, 67056 Ludwigshafen, Germany

6	Walther-Straub Institute of Pharmacology and Toxicology, Goethestrasse 33, 80336 Munich, Germany

7	VantAI, 151 W 42nd Street, New York, NY 10036, United States

8	NVIDIA DE GmbH, Einsteinstraße 172, 81677 München, Germany

9	Institute for Advanced Study (TUM-IAS), Lichtenbergstr. 2a, 85748 Garching/Munich, Germany

Table of Contents for SOM

### Short description of Supporting Online Material

In this document, we provide additional figures and their respective captions, supporting the *homo sapiens* case study, which is described in detail in **Results** of the main manuscript. The three figures (Fig. S1, S2, S3) show the visualization of 3D structures predicted by AlphaFold2 [1], and the respective membrane topology predicted by TMbed [2] for three proteins: Deoxyribonuclease (Dnase) I fragment (A0A192ZHB2), Macrophage-expressed gene 1 (Q2M385), and DnaJ homolog subfamily C member 11 (Q2M385).

All figures show the 3D protein structures with (**A**) a per-residue topology color-scheme: inside-to-outside TMH (light green), outside-to-inside TMH (dark green), inside-to-outside TMB (light blue), outside-to-inside TMB (dark blue), signal peptide (pink), other (grey), and (**B**) a per-residue AlphaFold2 color-scheme based on the confidence measure predicted local distance test (pLDDT): very low pLDDT ≤ 50 (red), low 50 < pLDDT ≤ 70 (yellow), confident 70 < pLDDT ≤ 90 (green), very confident pLDDT > 90 (blue).

### Material

Fig. S1: 3D structure and membrane topology visualization of a Deoxyribonuclease (Dnase) I fragment (A0A192ZHB2).

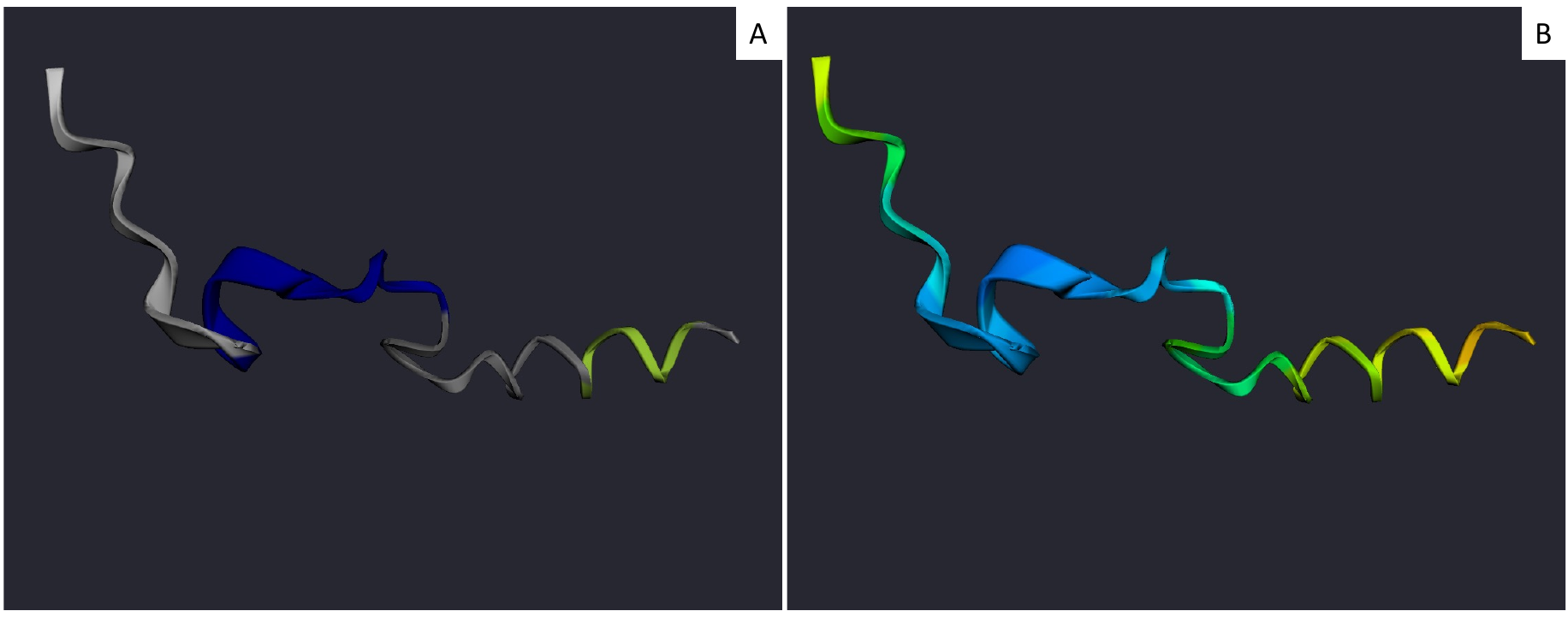

**Fig. S1: 3D structure and membrane topology visualization of a Deoxyribonuclease (Dnase) I fragment (A0A192ZHB2).** The visualization of the Dnase fragment in TMvis-DB is shown with transmembrane topology (TT) **(A)** and AlphaFold2 (AF2) **(B)** color-scheme as described in the introduction above. The two predicted TTs align with a helix and beta-strand predicted by AF2. The placement of the AF2 structure with both sections in a membrane seems unlikely and the average pLDDT is not confident. Further investigation is warranted to determine whether to evaluate the prediction of both TMbed and AF2.

Fig. S2: 3D structure and membrane topology visualization of the Macrophage-expressed gene 1 protein (Q2M385).

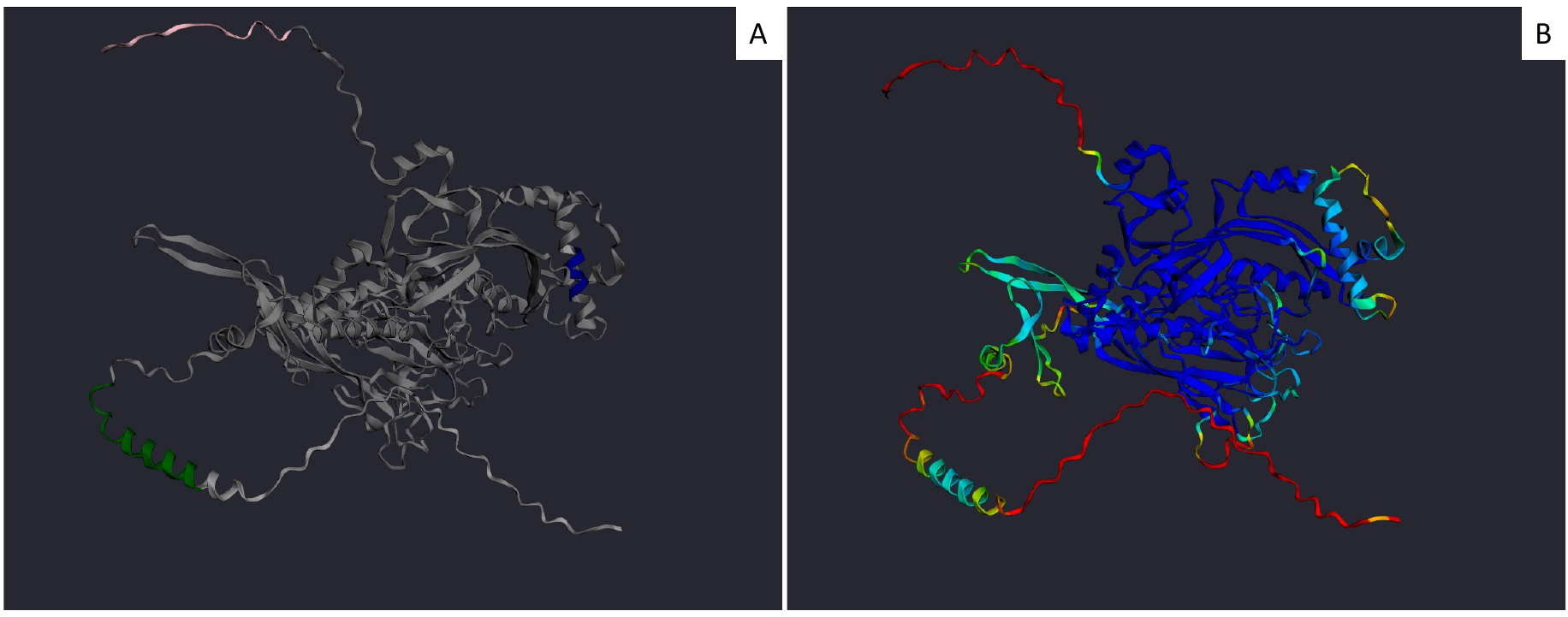

**Fig. S2: 3D structure and membrane topology visualization of the Macrophage-expressed gene 1 protein (Q2M385).** The protein is shown with TT **(A)** and AF2 **(B)** color-scheme as described in the introduction above. It has a predicted transmembrane helix (TMH) in a region of low to confident pLDDT that aligns with the predicted AF2 structure. The short predicted single transmembrane beta-strand (TMB) is likely a false positive as AF2 predicts an TMH with confident pLDDT.

Fig. S3: 3D structure and membrane topology visualization protein DnaJ homolog subfamily C member 11 (Q9NVH1).

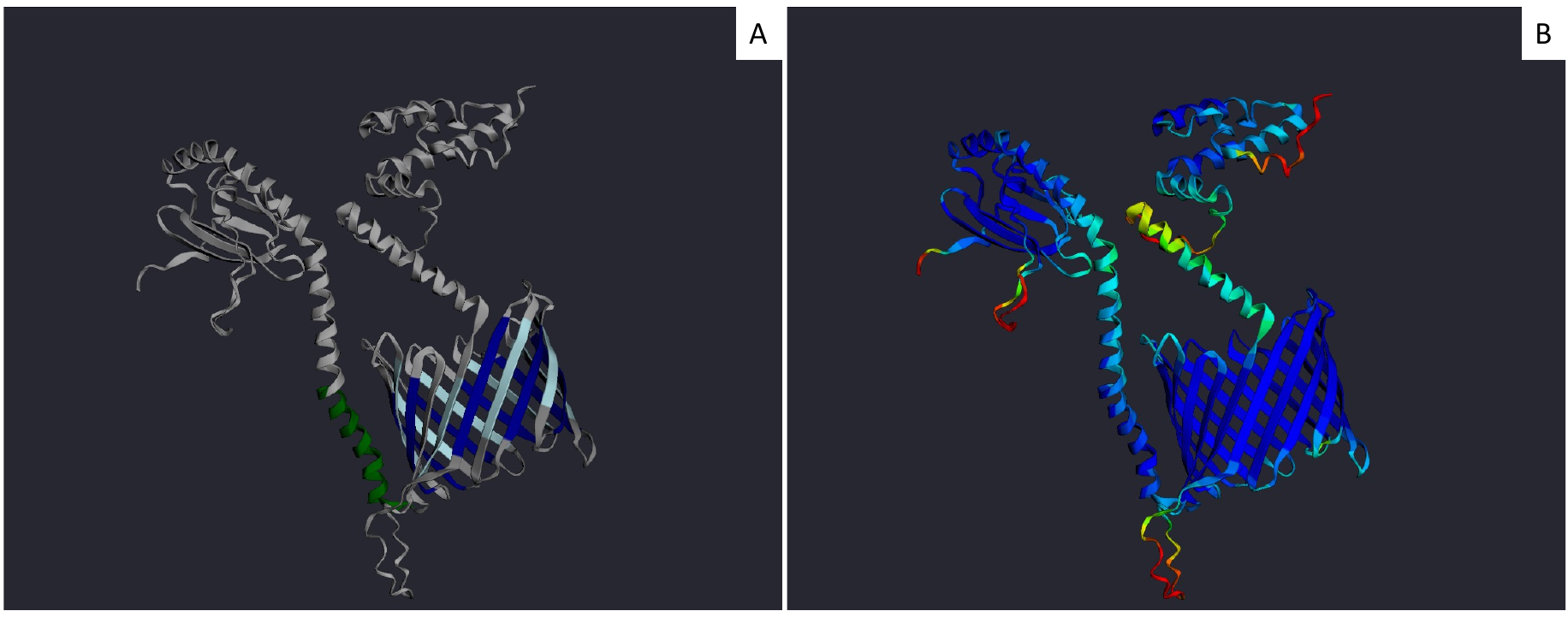

**Fig. S3: 3D structure and membrane topology visualization protein DnaJ homolog subfamily C member 11 (Q9NVH1).** The protein DnaJ with TT **(A)** and AF2 **(B)** color-scheme as described in the introduction above. The predicted TT aligns well with the predicted AF2 structure in regions of high pLDDT, and the length of the alpha-helix and beta-barrel could align with membrane boundaries.

### References for Supporting Online Material

[1]	J. Jumper, R. Evans, A. Pritzel, T. Green, M. Figurnov, O. Ronneberger, K. Tunyasuvunakool, R. Bates, A. Žídek, A. Potapenko, A. Bridgland, C. Meyer, S.A.A. Kohl, A.J. Ballard, A. Cowie, B. Romera-Paredes, S. Nikolov, R. Jain, J. Adler, T. Back, S. Petersen, D. Reiman, E. Clancy, M. Zielinski, M. Steinegger, M. Pacholska, T. Berghammer, S. Bodenstein, D. Silver, O. Vinyals, A.W. Senior, K. Kavukcuoglu, P. Kohli, D. Hassabis, Highly accurate protein structure prediction with AlphaFold, Nature. 596 (2021) 583–589. https://doi.org/10.1038/s41586-021-03819-2.

[2]	M. Bernhofer, B. Rost, TMbed: transmembrane proteins predicted through language model embeddings, BMC Bioinformatics. 23 (2022) 326. https://doi.org/10.1186/s12859-022-04873-x.
